# Vagal interoception of microbial metabolites from the small intestinal lumen

**DOI:** 10.1101/2023.12.18.572257

**Authors:** Kelly G. Jameson, Sabeen A. Kazmi, Celine Son, Donya Mazdeyasnan, Emma Leshan, Helen E. Vuong, Jorge Paramo, Arlene Lopez-Romero, Long Yang, Felix E. Schweizer, Elaine Y. Hsiao

## Abstract

The vagus nerve is proposed to enable communication between the gut microbiome and brain, but activity-based evidence is lacking. Herein, we assess the extent of gut microbial influences on afferent vagal activity and metabolite signaling mechanisms involved. We find that mice reared without microbiota (germ-free, GF) exhibit decreased vagal afferent tone relative to conventionally colonized mice (specific pathogen-free, SPF), which is reversed by colonization with SPF microbiota. Perfusing non-absorbable antibiotics (ABX) into the small intestine of SPF mice, but not GF mice, acutely decreases vagal activity, which is restored upon re-perfusion with bulk lumenal contents or sterile filtrates from the small intestine and cecum of SPF, but not GF, mice. Of several candidates identified by metabolomic profiling, microbiome-dependent short-chain fatty acids, bile acids, and 3-indoxyl sulfate stimulate vagal activity with varied response kinetics, which is blocked by co-perfusion of pharmacological antagonists of FFAR2, TGR5, and TRPA1, respectively, into the small intestine. At the single-unit level, serial perfusion of each metabolite class elicits more singly responsive neurons than dually responsive neurons, suggesting distinct neuronal detection of different microbiome- and macronutrient-dependent metabolites. Finally, microbial metabolite-induced increases in vagal activity correspond with activation of neurons in the nucleus of the solitary tract, which is also blocked by co-administration of their respective receptor antagonists. Results from this study reveal that the gut microbiome regulates select metabolites in the intestinal lumen that differentially activate chemosensory vagal afferent neurons, thereby enabling microbial modulation of interoceptive signals for gut-brain communication.

**HIGHLIGHTS:**

- Microbiota colonization status modulates afferent vagal nerve activity
- Gut microbes differentially regulate metabolites in the small intestine and cecum
- Select microbial metabolites stimulate vagal afferents with varied response kinetics
- Select microbial metabolites activate vagal afferent neurons and brainstem neurons via receptor-dependent signaling

## INTRODUCTION

The gut microbiota is emerging as a key modulator of brain function and behavior, as several recent studies reveal effects of gut microbes on neurophysiology, complex animal behaviors, and endophenotypes of neurodevelopmental, neurological, and neurodegenerative diseases.^1-3^ Despite these findings supporting a “microbiome-gut-brain axis”, the mechanisms that underlie interactions between gut microbes and the brain remain poorly understood. While many studies highlight neuroimmune pathways for microbial influences on the brain,^4,5^ it is also believed that the gut microbiota may directly signal to the brain via gut-innervating vagal neurons.^6^ However, existing evidence for the vagal route largely derives from studies wherein subdiaphragmatic vagotomy abrogates behavioral alterations in response to microbiota perturbation in mice.^2,7-10^ This ablates signaling in both afferent and efferent directions, not only to the intestine, but also with other peripheral organs. While the approach provides an important initial indication that the vagus nerve contributes to behavioral phenotypes that are modified by the microbiome, *in vivo* evidence of microbial signaling through vagal neurons is needed, and fundamental questions remain regarding the nature of microbial effects on vagal activity, the particular molecular constituents involved, and the diversity of neuronal responses elicited.

The gut microbiome is central to dietary metabolism and modulates hundreds of biochemicals in the intestine, as well as the blood and various distal extraintestinal tissues.^11,12^ Biochemical screens of supernatants from cultured human-derived gut microbes find that soluble microbial products (largely uncharacterized) have the capacity to directly bind to numerous G-protein-coupled receptors (GPCRs) that mediate neurotransmitter and neuropeptide signaling,^13,14^ some of which are reportedly expressed by vagal neurons.^15-19^ As such, microbial metabolites generated in the intestinal lumen have the potential to directly or indirectly activate vagal neurons via receptors on mucosal sensory afferents, or on enteroendocrine cells that synapse onto the mucosal sensory afferents, as has been described for select lumenal nutrient stimuli^20-22^ and microbial antigens.^23,24^ In this study, we assess afferent vagal nerve activity in response to the presence, absence, depletion, and restoration of the gut microbiota. We further profile microbiome-dependent metabolites within the proximal small intestine and cecum, which are poised to signal to gut mucosal vagal afferents. We identify select microbial metabolites that induce vagal afferent neuronal activity with varied kinetics when administered to the lumen of the small intestine. Finally, we apply pharmacological approaches to probe the potential for direct and/or indirect receptor-mediated signaling of microbial metabolites to vagal neurons. This study provides fundamental functional and mechanistic evaluation of the gut microbiome as a regulator of lumenal metabolites that modulate vagal chemosensory signaling across the gut-brain axis.

## RESULTS

### The gut microbiome promotes afferent vagal nerve activity

To determine the extent to which the microbiome contributes to gut-brain signaling via the vagus nerve, we began by applying whole nerve electrophysiology to measure bulk activity of vagal afferents in wildtype C57BL/6J mice reared in the presence vs. absence of microbial colonization (**Figure 1A, left**). Germ-free (GF) mice exhibited significantly decreased afferent vagal nerve activity as compared to conventionally colonized (specific pathogen-free, SPF) controls (**Figure 1B-1C**). These reductions were reversed by colonizing GF mice with the SPF microbiota during adulthood (conventionalized, CONV), suggesting active interactions between the microbiome and vagus nerve that occur independently of developmental colonization. Treating adult SPF mice with oral broad-spectrum antibiotics (ABX; ampicillin, neomycin, vancomycin, and metronidazole) for 7 days to reduce bacterial load yielded modest, but not statistically significant, reductions in afferent vagal nerve activity. We hypothesized that the inconsistent phenotype between the ABX and GF conditions may be due to incomplete depletion of bacteria by ABX treatment, enrichment of native ABX-resistant bacteria over the one-week treatment period,^25^ off-target effects of ABX absorbed into the systemic circulation,^26^ the activity of non-bacterial members of the microbiota,^27^ and/or confounding effects of developmental GF rearing.^28^ To gain further insight, we recorded afferent vagal nerve activity acutely while introducing a constant flow of the subset of the ABX cocktail that is non-absorbable (i.e., vancomycin and neomycin) directly into the lumen of the duodenum^29^ and through the first ∼10 cm of the small intestine, a site of dense vagal innervation^16^ (**Figure 1A, right**). Perfusing nonabsorbable ABX into the intestinal lumen of SPF mice decreased afferent vagal nerve activity, as compared to vehicle (VEH)-perfused controls (**Figure 1D-1F**). ABX-induced decreases in afferent vagal nerve activity were not seen in GF mice perfused with ABX (**Figure 1D-1F**). These results suggest that intestinal perfusion with non-absorbable ABX decreases afferent vagal nerve activity via the bactericidal actions of ABX on microbes in the small intestine.

**Figure 1:**
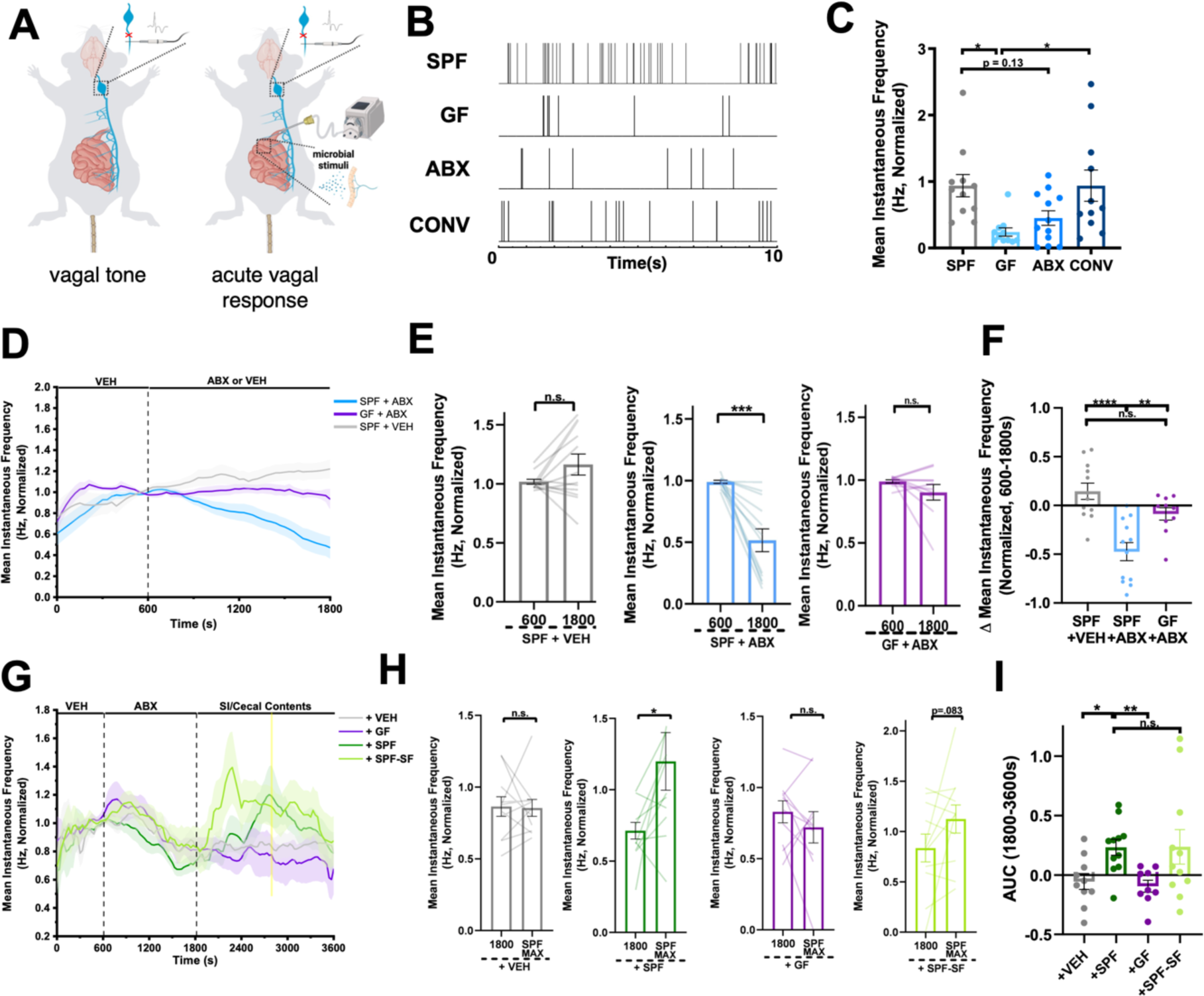
The Gut Microbiota and Lumenal Microbial Metabolites Promote Vagal Afferent Activity. **A**) Diagram of *in vivo* whole nerve vagal electrophysiology for quantification of vagal tone (left, for data in B-C) or acute afferent vagal nerve response to lumenal stimuli (right, for data in D-I) **B**) Representative images of detected vagal spikes (vertical lines) over 10s in conventionally colonized (SPF), germ-free (GF), antibiotic-treated (ABX), and conventionalized GF (CONV) mice. **C**) Vagal afferent tone of SPF, GF, ABX, and CONV mice. (SPF, n = 11 mice; GF, n = 11 mice; ABX, n = 12 mice; CONV, n = 11 mice). One-Way ANOVA + Tukey. **D**) Afferent vagal nerve response in SPF or GF mice perfused intestinally with non-absorbable antibiotics (ABX, vancomycin/neomycin, 1mg/mL) or vehicle (VEH) (SPF +VEH, n = 12 mice; SPF +ABX, n = 12 mice; GF + ABX, n = 10 mice). **E**) Afferent vagal nerve firing rates before and after intestinal perfusion of SPF or GF mice with ABX or VEH. (SPF +VEH, n = 12 mice; SPF +ABX, n = 12 mice; GF + ABX, n = 10 mice). Paired t-test. **F**) Change in afferent vagal nerve activity after intestinal perfusion with ABX or VEH (t=1800) relative to stable baseline (t=600s) (SPF +VEH, n = 12 mice; SPF +ABX, n = 12 mice; GF + ABX, n = 10 mice). One-way ANOVA + Tukey. **G**) Afferent vagal nerve response in mice intestinally perfused with ABX, followed by re-perfusion with pooled small intestinal (SI) and cecal contents from SPF mice (+SPF) or GF mice (+GF), sterile filtered SI/cecal contents from SPF mice (+SPF-SF), or VEH (+VEH, n = 10 mice; +SPF, n = 11 mice; +GF, n = 9 mice; +SPF-SF, n = 11 mice). **H**) Afferent vagal nerve firing rate after intestinal ABX perfusion (t=1800s) and after re-perfusion with VEH, SI/cecal contents from SPF mice, SI/cecal contents from GF mice, or sterile filtered SI/cecal contents from SPF mice, at the time of maximum mean firing rate for perfusion of SPF SI/Cecal contents (SPF MAX, t = 2760s) (+VEH, n = 10 mice; +SPF, n = 11 mice; +GF, n = 9 mice; +SPF-SF, n = 11 mice). Paired t-test. **I**) Afferent vagal nerve activity as measured by area under the curve (AUC) (from 1800-3600s) in response to intestinal perfusion with VEH (n = 10 mice), SPF SI/cecal contents (n = 11 mice), GF SI/cecal contents (n = 9 mice), and SPF-SF SI/cecal contents (n = 11 mice). Brown-Forsythe and Welch ANOVA + Games-Howell. All data displayed as mean +/- SEM, *p < 0.05, **p < 0.01, ***p < 0.001, **** p < 0.0001.

The gut microbiota influences many aspects host biology, in large part by bacterial metabolism of dietary substrates, synthesis of secondary metabolites, and modification of host-derived molecules in the intestine.^4,30-33^ To evaluate the effects of lumenal microbial molecules on afferent vagal nerve activity, we administered a solution of small intestinal (SI) and cecal contents collected from donor SPF or GF mice into the SI lumen of ABX-perfused SPF mice. SI perfusion with non-absorbable ABX reduced afferent vagal nerve activity, as reported above, whereas re-perfusion of SI/cecal contents from SPF mice acutely increased activity toward levels seen at pre-ABX baselines (“+ SPF” in **Figure 1G-1I**). No such effect was seen with re-perfusion of SI/cecal contents from GF mice (“+ GF” in **Figure 1G-1I**), suggesting that the observed vagal response is due the presence of SI and cecal microbes and/or microbial molecules. To investigate the contribution of microbial small molecules, in particular, we sterile-filtered the equivalent solution of SPF SI/cecal contents and administered it to the SI lumen of ABX-perfused SPF mice. Sterile-filtered SPF SI/cecal contents increased afferent vagal nerve activity to levels similar to SPF on average, albeit with shorter latency and more variability (“+ SPF-SF” in **Figure 1G-1I**). These differences could be due to differential kinetics of small molecule signaling resulting from a lack of macromolecules present in SPF-SF samples^20^ and/or variability in the fidelity of small molecules upon filtration or the distribution of small-molecule activated receptors along the length of the proximal and medial small intestine.^34^ Taken together, these data provide evidence that active signaling from the gut microbiota modulates vagal afferent activity *in vivo*, and that these effects are mediated, at least in part, by microbial small molecules within the lumen of the small intestine and cecum.

### Microbiome-dependent bile acids, short-chain fatty acids, and 3-indoxyl sulfate stimulate afferent vagal nerve activity in a receptor-dependent manner

The gut microbiota regulates numerous metabolites within the host.^11,12^ However, most characterizations to date have profiled metabolites in fecal or serum samples, excluding signaling molecules localized to the small intestine and cecum that are poised to interact with villus-innervating vagal neurons.^35^ To identify candidate microbial metabolites in the small intestine and cecum that may modify vagal afferent activity, we performed liquid chromatography-tandem mass spectrometry (LC-MS/MS) based untargeted metabolomic profiling of lumenal contents from the duodenum (proximal SI) and cecum of SPF, GF, ABX, and CONV mice. 931 metabolites were identified from mouse proximal SI and cecal contents (**Tables S1 and S2**). Principal component analysis revealed distinct clustering of GF and ABX samples away from SPF and CONV samples along PC1 (**Figure 2A**), indicating that acute ABX depletion of the gut microbiota yields SI and cecal metabolomic profiles that are similar to those seen with GF rearing and that adult inoculation of GF mice with a conventional microbiota induces SI and cecal metabolomic profiles that are similar to those seen with conventional colonization (SPF). This is consistent with our finding that acute depletion and re-introduction of the microbiota or microbial metabolites alters afferent vagal nerve activity (**Figure 1**). However, there were also notable differences within the cecal datasets in particular, with discrimination of ABX from GF profiles and, to a lesser degree, CONV from SPF profiles, along PC2 (**Figure 2A**, bottom). These differences highlight potential developmental influences of microbial colonization on host physiology^36^ and/or incomplete depletion and/or restoration of microbial communities within the lower GI tract relative to the proximal small intestine.^37^ Based on the ability of both GF status and acute perfusion of nonabsorbable ABX to decrease afferent vagal nerve activity and of CONV to elevate activity toward levels seen in SPF controls (**Figure 1A-1F**), we then filtered the datasets to identify metabolites that were commonly differentially regulated by both microbiota-deficient conditions relative to both colonized conditions. In SI contents, there were 79 shared metabolites that were significantly modulated by both GF and ABX conditions relative to both SPF and CONV conditions, and in cecal contents, there were 521 (**Figure 2B, Table S1 and S2**). Based on the observation that SI/cecal contents and filtrates from SPF mice stimulate afferent vagal nerve activity compared to filtrates from GF controls (**Figure 1G-1I**), we focused in particular on the subset of 49 SI and 335 cecal metabolites that were significantly decreased by microbiota deficiency relative to colonized conditions (**Figure 2B**, **Table S1 and S2**). These included microbial metabolites that were extremely low or undetectable in microbiota-deficient conditions (which we refer to as “microbiome-dependent”), as well as metabolites that were partially downregulated (but still detectable) in microbiota-deficient conditions (which we refer to as “microbially modulated”).

**Figure 2:**
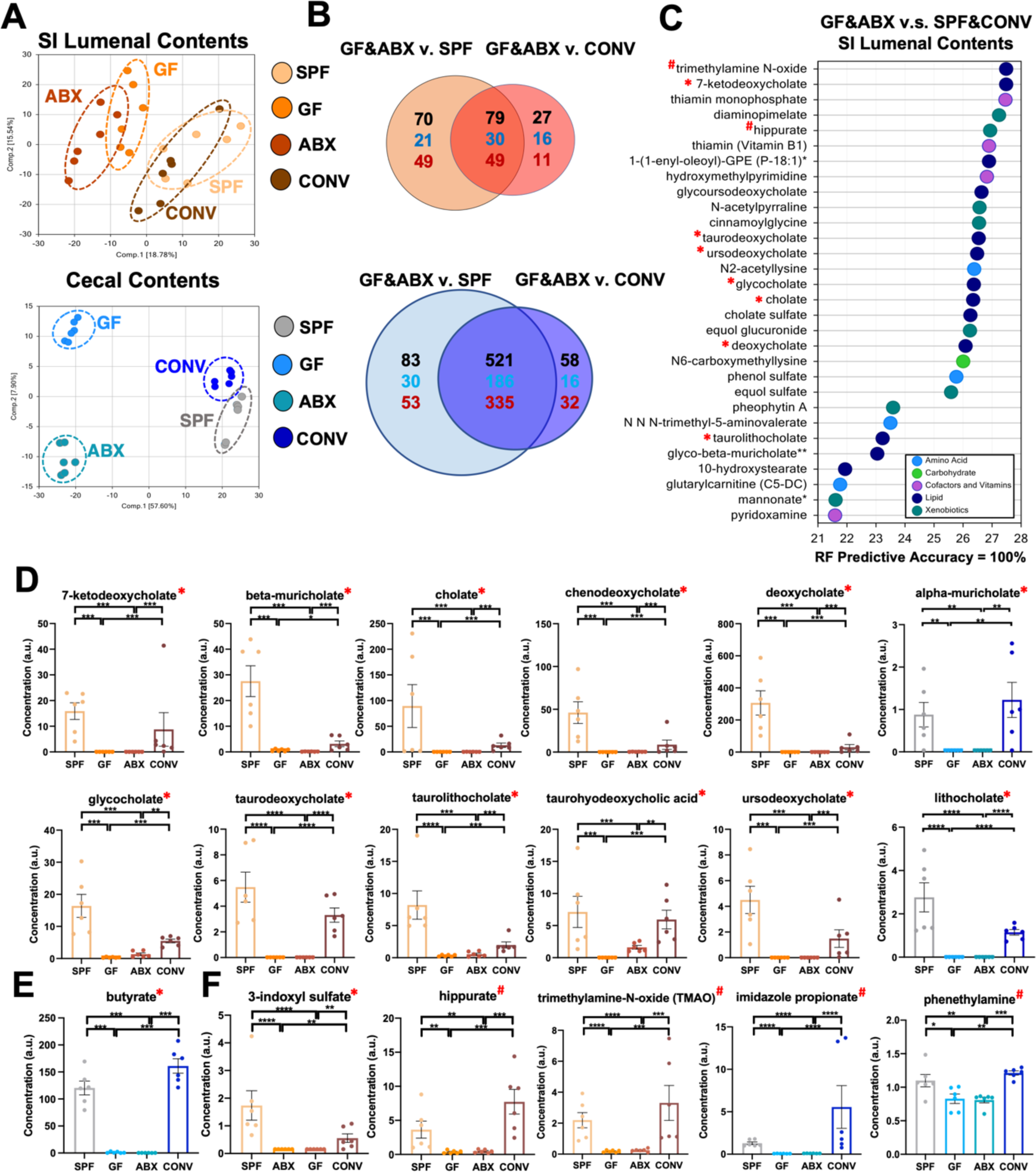
The Gut Microbiome Regulates Metabolites in the Lumen of the Small Intestine and Cecum. **A**) PCA analysis of metabolomic data from small intestinal (SI, top) or cecal (bottom) lumenal contents from SPF, GF, ABX, and CONV mice. (n=6 mice for all groups). **B**) Venn diagram of differentially modulated metabolites in SI (top) and cecal (bottom) lumenal contents. Red numbers indicate downregulated metabolites and blue numbers indicate upregulated metabolites. (n=6 mice for all groups). **C**) Random forest (RF) analysis of metabolomic data from SI lumenal contents reveals the top 30 metabolites that distinguish GF/ABX from SPF/CONV samples with 100% predictive accuracy. Red asterisks indicate metabolites included in experiments in Figure 3. Hash symbols indicate metabolites included in experiments in Supplemental Figure 2 (n=6 mice for all groups). **D-E**) Lumenal levels of microbially modulated bile acids and the short-chain fatty acid butyrate from SI (orange tones) and/or cecum (blue tones) of SPF, GF, ABX, and CONV mice. (n=6 mice for all groups). Welch’s t-test. **F**) Lumenal levels of microbially modulated metabolites with unknown signaling to vagal neurons from SI (orange tones) or cecum (blue tones) of SPF, GF, ABX, and CONV mice. (n=6 mice for all groups). Welch’s t-test. All data displayed as mean +/- SEM, *p < 0.05, **p < 0.01, ***p < 0.001, ****p<.0001

To identify the subset of microbial metabolites that have the potential to signal directly to vagal neurons, we cross-referenced existing bulk and single-cell RNA sequencing datasets^15,16,18,19,38^ for reported expression of known or putative receptors for the candidate SI and cecal metabolites. This identified select species of microbiome-dependent bile acids (BAs), a subset of which were identified as key drivers for classifying microbiota status via random-forest analysis in small-intestinal samples (**Figure 2C-2D**). Additional classes of metabolites that were uncovered included the microbiome-dependent short chain fatty acid (SCFA) butyrate (**Figure 2E**) and microbially modulated tryptophan derivatives (TRPs, **Figure S1A**), fatty acid ethanolamides (FAEs, **Figure S1B**), monohydroxy fatty acids (MFAs, **Figure S1C**), as well as succinate and glutamate (**Figure S1D-S1E**). To initially assess the ability of these metabolites to acutely modify vagal activity from the SI lumen, we recorded afferent vagal nerve activity while perfusing physiologically relevant concentrations of metabolite pools into the SI. There were no statistically significant changes in afferent vagal nerve activity with SI perfusion of the detected TRPs, FAEs, MFAs, or succinate (**Figure S2A-S2D**). Consistent with existing literature demonstrating vagal responses to gastric delivery of glutamate,^39^ SI perfusion of glutamate robustly increased afferent vagal nerve activity (**Figure S2E**). We further observed that SI perfusion of select microbiome-dependent BAs elicited rapid, transient afferent vagal nerve activity (**Figure 3A-3C**), which parallels existing literature on systemic administration of select primary and secondary BAs.^31^ In contrast, SI perfusion of microbiome-dependent SCFAs (acetate, propionate, and butyrate) led to slower onset and gradual increases in afferent vagal nerve activity (**Figure 3D-3F**). This latency could be due to metabolite-specific differences in the rate of intestinal absorption,^40^ differential spatial localization of metabolite absorption and functional activity along the length of the gastrointestinal tract,^41,42^ and/or indirect signaling of the metabolites to non-neuronal mediators.^21,43,44^

**Figure 3:**
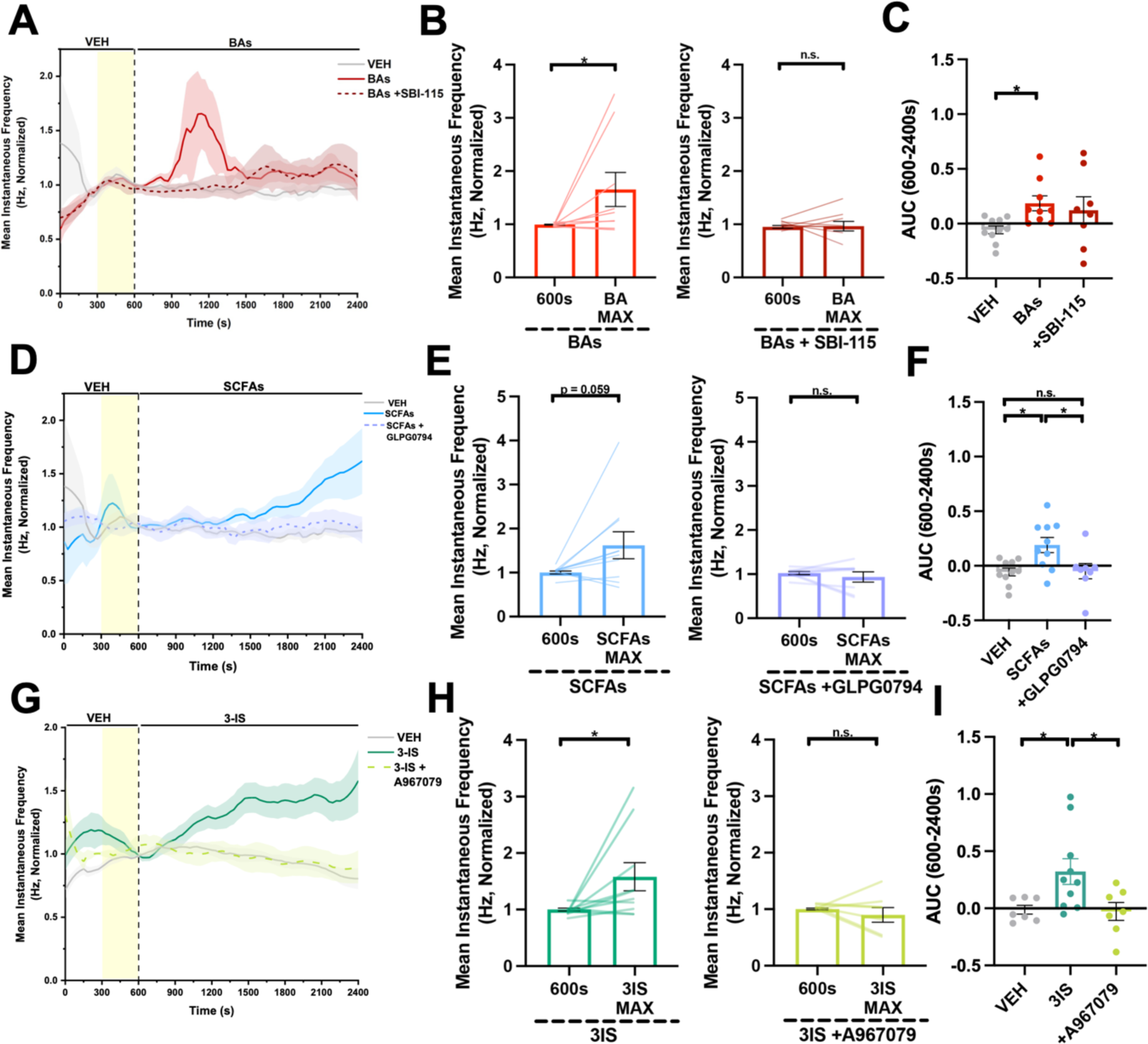
Select Lumenal Microbial Metabolites Increase Afferent Vagal Nerve Activity with Varied Response Kinetics. **A**) Afferent vagal nerve firing rate in SPF mice after intestinal perfusion with vehicle (VEH: PBS, n=10 mice) or pooled bile acids (BAs: cholate, 1240nM; glycocholate, 3.5nM; chenodeoxycholate, 42nM; alpha-muricholate, 142nM; beta-muricholate, 1080nM; deoxycholate, 390nM; taurodeoxycholate, 260nM; ursodeoxycholate, 74nM; taurohyodeoxycholate, 18.8nM; 7-ketodeoxycholate, 100nM; lithocholate, 390nM; taurolithocholate, 0.33nM; n = 9 mice), with or without pre- and co-perfusion with TGR5 antagonist (SBI-115, 200uM, n = 8 mice). Yellow shading indicates perfusion of pure antagonist prior to co-perfusion of antagonist with metabolites. **B**) Normalized afferent vagal nerve firing rate before and during treatment with BAs (left, n = 9 mice), or BAs and SBI-115 (right, n = 8 mice). BA max, t = 1114s. Wilcoxon matched-pairs signed rank test. **C**) Afferent vagal nerve firing rate as measured by area under the curve (AUC) (from 600-2400s) in response to intestinal perfusion with VEH (PBS, n = 10 mice) or pooled BAs (n = 9 mice), with or without pre- and co-perfusion with TGR5 antagonist (SBI-115, 200uM, n = 8 mice). Brown-Forsythe and Welch ANOVA + Games-Howell. **D**) Afferent vagal nerve firing rate in SPF mice after intestinal perfusion with VEH (PBS, n = 10 mice) or pooled short-chain fatty acids (SCFAs: acetate, 80uM; butyrate, 22uM; propionate, 10uM, pooled, n = 10 mice), with or without pre- and co-perfusion with FFAR2 antagonist (GLPG0794, 10uM, n = 8 mice). Yellow shading indicates perfusion of pure antagonist prior to co-perfusion of antagonist with metabolites. **E**) Normalized afferent vagal nerve firing rate before and during treatment with SCFAs (left, n = 10 mice), or SCFAs and GLPG0794 (right, n = 8 mice). SCFA max, t = 2400s. Paired t-test. **F**) Afferent vagal nerve firing rate as measured by AUC (from 600-2400s) in response to intestinal perfusion with VEH (PBS, n = 10 mice) or pooled SCFAs (10uM, n = 10 mice), with or without pre- and co-perfusion with FFAR2 antagonist (GLPG0794, 10uM, n = 8 mice). One-way ANOVA + Tukey. **G**) Afferent vagal nerve firing rate in SPF mice after intestinal perfusion with VEH (1uM KCl, n = 7 mice) or 3-indoxyl sulfate (3-IS, 1uM, n = 10 mice), with or without pre- and co-perfusion with TRPA1 antagonist (A967079, 10uM, n = 7 mice). Yellow shading indicates perfusion of pure antagonist prior to co-perfusion of antagonist with metabolites. **H**) Normalized afferent vagal nerve firing rate before and during treatment with 3IS (left, n = 10 mice), or 3IS and A967079 (right, n = 7 mice). 3IS max, t = 2400s. Paired t-test. I) Afferent vagal activity as measured by AUC (from 600-2400s) in response to intestinal perfusion with VEH (1uM KCl, n=7 mice) or 3-indoxyl sulfate (3-IS, 1uM, n = 10 mice), with or without pre- and co-perfusion with TRPA1 antagonist (A967079, 10uM, n = 7 mice). One-way ANOVA + Tukey. All data displayed as mean +/- SEM, *p < 0.05.

Microbiome-dependent BAs and SCFAs in the intestinal lumen have the potential to bind to cognate receptors expressed by various cell types in the gastrointestinal tract (e.g., vagal, enteroendocrine, epithelial, immune)^45,46^. To determine the relative contributions of different cognate GPCRs to vagal responses induced by lumenal microbial metabolites, we perfused select receptor antagonists immediately before and during administration of their corresponding metabolites into the SI lumen of SPF mice. BAs signal through the membrane-bound Takeda G protein-coupled receptor 5 (TGR5), which is expressed by gut-innervating vagal neurons^31^ as well as intestinal epithelial cells and various intestinal innate immune cells.^32,45,47^ Intestinal pre- and co-perfusion of the TGR5 antagonist m-tolyl 5-chloro-2-[ethylsulonyl] pyrimidine-4-carboxylate (SBI-115)^48^ prevented the initial rapid, transient increases in afferent vagal nerve activity induced by microbiome-dependent BAs (**Figure 3A-3C**). We do not observe a difference in total area under the curve (AUC) across the entire stimulus window, as administration of SBI-115 leads to a delayed rise in vagal activity that was not observed with perfusion of microbiome-dependent BAs alone. This suggests that TGR5 antagonism may elicit compensatory vagal responses to microbiome-dependent BAs through farsenoid X receptor (FXR) or other signaling mechanisms independent of local TGR5. SCFAs signal to free fatty acid receptor 2 (FFAR2), which is expressed by intestinal epithelial cells^41^ and free fatty acid receptor 3 (FFAR3), which is expressed by gut-innervating vagal neurons.^49^ Intestinal pre- and co-perfusion of the FFAR2 antagonist 4- [[(R)-1-(benzo[b]thiophene-3-carbonyl)-2-methyl-azetidine-2-carbonyl]-(3-chloro-benzyl)-amino]- butyric acid 99 (GLPG0974)^50^ prevented the increase in afferent vagal nerve activity induced by SCFAs (**Figure 3D-3F**). These results suggest that microbiome-dependent SCFAs likely elevate vagal activity via indirect activation of intestinal epithelial cells or other cellular mediators.

In addition to testing microbial metabolites with reported receptor expression by vagal neurons, we also evaluated effects of select microbiome-dependent metabolites that have as yet unknown signaling mechanisms on vagal activity. We focused in particular on metabolites that i) are reproducibly dependent upon the microbiome across various studies and biological contexts^51^ and ii) have been reportedly linked to brain function and/or behavior.^1,52-55^ Of these, 3-indoxyl sulfate (3IS), hippurate, and trimethylamine-N-oxide (TMAO) are microbiome-dependent metabolites in SI lumen, imidazole propionate is a microbiome-dependent metabolite in the cecum, and phenethylamine is microbially modulated in the cecum (**Figure 2F**). These metabolites are also reduced in the serum of microbiome-deficient mice,^11^ suggesting that they are typically absorbed from the intestine and poised to interact with mucosal vagal afferents. Perfusing physiologically relevant concentrations of hippurate, TMAO, imidazole propionate, and phenethylamine individually through the small intestine had no measurable effect on afferent vagal nerve activity (**Figure S2F-S2I**). In contrast, SI perfusion with 3IS elicited rapid and sustained increases in afferent vagal nerve activity relative to vehicle controls (**Figure 3G-3I**). Microbial indole (the metabolic precursor to 3IS) and its derivative indole-3-carboxaldehyde are reported to activate vagal afferent neurons in zebrafish via indirect stimulation of colonic enteroendocrine cells in a transient receptor potential ankyrin1 (TRPA1)-dependent manner.^43^ To evaluate this possible signaling mechanism for 3IS in the small intestine, we pre- and co-perfused the TRPA1 antagonist (1E,3E)-1-(4-Fluorophenyl)-2-methyl-1-penten-3-one oxime (A967079)) with 3IS into the SI lumen, which completely prevented 3IS-induced afferent vagal nerve activity (**Figure 3G-3I**). Overall, these results reveal that microbial BAs, SCFAs, and 3IS promote afferent vagal nerve activity through receptor-dependent signaling from the SI lumen.

### Lumenal BAs, SCFAs, and 3IS excite both distinct and shared subsets of afferent vagal neurons with varied temporal responses

Different lumenal stimuli can activate distinct populations of vagal neurons with differing response kinetics. In particular, recent work has identified populations of afferent vagal neurons that respond exclusively to fats versus sugars.^22^ Microbiome-dependent BAs, SCFAs, and 3IS are related to dietary metabolism of fats, complex carbohydrates, and proteins, respectively,^30,43,56^ raising the question of whether they promote vagal nerve activity via shared vs. distinct vagal afferent neurons. To gain insight, we imaged calcium activity of vagal afferent neurons in response to acute SI perfusion of microbiome-dependent metabolites in mice expressing GCaMP6s in Phox2b+ sensory neurons (**Figure 4A-B**). Microbiome-dependent BAs, SCFAs, and 3IS elicited calcium responses in 57%, 48%, and 58% of detected vagal afferent neurons, respectively, on average across independent animals (**Figure 4C**). The latency to maximum calcium response varied within each subclass of microbial metabolite, where SCFAs and 3IS similarly elicited primarily delayed calcium responses, while BAs elicited a bimodal distribution of acute and delayed calcium responses (**Figure 4D**). Upon perfusing pairs of metabolites in counterbalanced sequence, BAs and SCFAs elicited calcium responses in largely distinct vagal afferent neurons, with 43% responsive to BAs only, 38% responsive to SCFAs, and 19% dually responsive to both BAs and SCFAs (“BAs <> SCFAs” in **Figure 4E**). In contrast, sequential perfusion of 3IS and SCFAs yielded many shared neuronal responses, where 40% of vagal afferent neurons were dually responsive to 3IS and SCFAs, 27% to SCFAs only, and 33% to 3IS only (“3IS <> SCFAs” in **Figure 4E)**. Similarly, with BAs and 3IS, we observed 43% dual responders, 35% responsive to 3IS only, and 22% responsive to BAs only (“BAs <> 3IS” in **Figure 4E**). Together, these data reveal distinct and shared neuronal populations for sensing different microbiome- and macronutrient-dependent metabolites, with greater distinct neuronal responses to microbial BAs and SCFAs, than either with 3IS.

**Figure 4:**
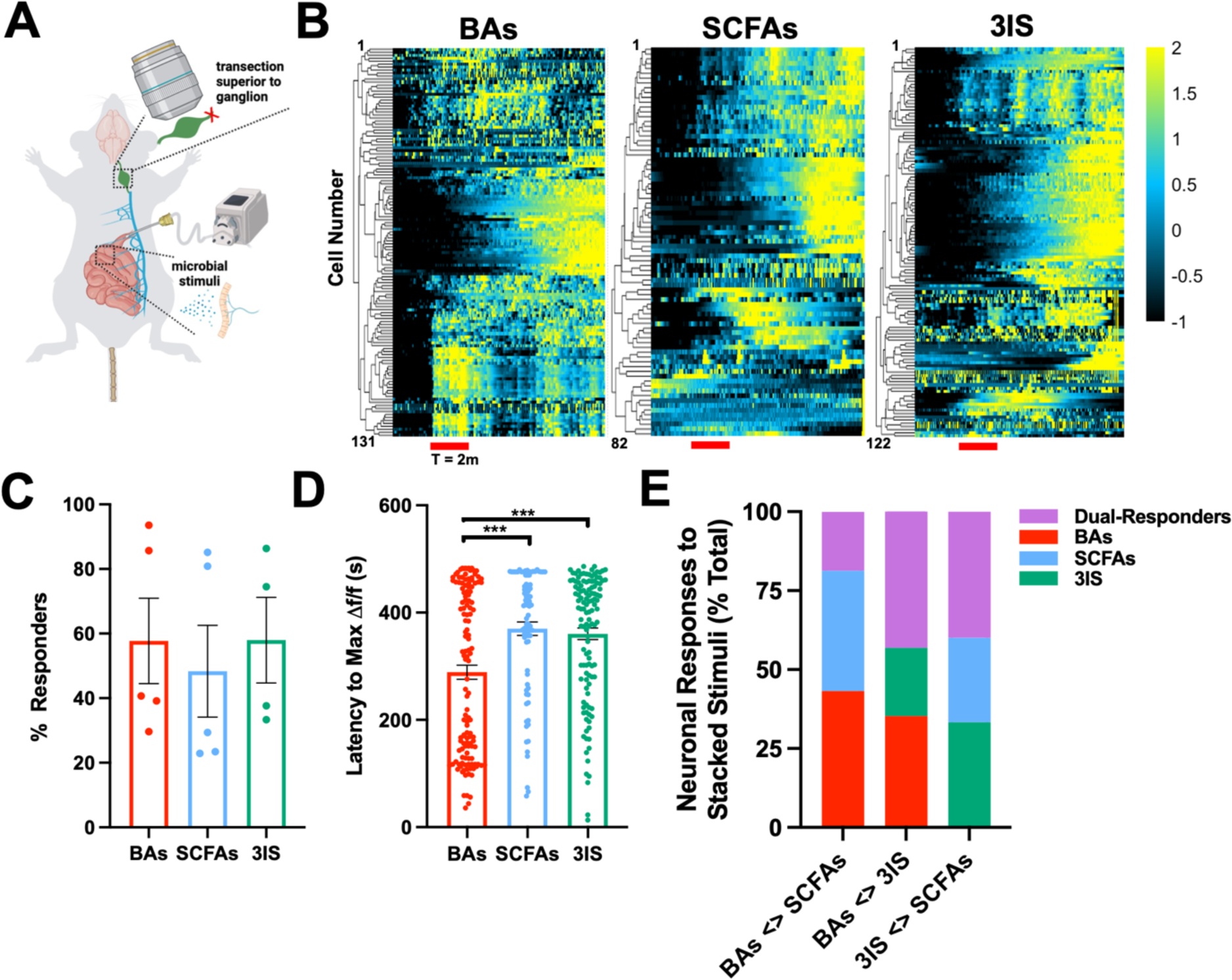
Lumenal BAs, SCFAs, and 3IS activate distinct subsets of vagal afferent neurons with heterogeneous kinetics. **A**) Diagram of experimental setup for *in vivo* calcium imaging. **B**) Representative heatmaps for cells responding to only pooled bile acids (BAs: cholate, 1240nM; glycocholate, 3.5nM; chenodeoxycholate, 42nM; alpha-muricholate, 142nM; beta-muricholate, 1080nM; deoxycholate, 390nM; taurodeoxycholate, 260nM; ursodeoxycholate, 74nM; taurohyodeoxycholate, 18.8nM; 7-ketodeoxycholate, 100nM; lithocholate, 390nM; taurolithocholate, 0.33nM; n = 4 mice, n = 131 units, right), only pooled short-chain fatty acids (SCFAs: acetate, 80uM; butyrate, 22uM; propionate, 10uM, n = 4 mice, n = 82 cells, middle), or only 3-indoxyl sulfate (3IS, 1uM, n = 4 mice, n = 132 cells, left). Recording duration for all experiments was 10 minutes. **C**) Percentage of metabolite-responsive neurons out of total excitable neurons in response to lumenal perfusion of BAs (n = 5 mice), SCFAs (n = 5 mice), or 3IS (n = 4 mice). **D**) Latency to maximum change in fluorescence for metabolite-responding neurons with lumenal perfusion of BAs (n = 4 mice), SCFAs (n = 4 mice) or 3IS (n = 4 mice). One-way ANOVA + Tukey. **E**) Percentage of single- or dual-responding neurons following serial perfusion of BAs (n = 5 mice), SCFAs (n = 5 mice), and 3IS (n = 4 mice). Order of metabolites for perfusion was counterbalanced between experiments. All data displayed as mean +/- SEM, ***p < 0.001

### Receptor-mediated signaling of BAs, SCFAs, and 3IS from the small intestinal lumen activates neurons in the NTS

Changes in the gut microbiota have been associated with altered activation of neurons of the nucleus of the solitary tract (NTS), which receives direct visceral afferents from nodose neurons.^57,58^ Consistent with the ability of microbial metabolites in the SI lumen to stimulate vagal afferent neuronal activity (**Figures 3 and 4**), we find that acute lumenal perfusion of microbiome-dependent BAs, SCFAs, and 3IS each increase neuronal expression of the activation marker cFos in the NTS (**Figure 5A-B**), to levels similar to those seen with intestinal perfusion of sucrose.^21,59^ As with the afferent vagal nerve and neuronal responses, the microbial metabolite-driven increases in NTS neuronal activation were prevented by pre- and co-administration of antagonists for TGR5, FFAR2, and TRPA1 with their respective metabolite ligands. Together, these data indicate that microbial BAs, SCFAs, and 3IS in the SI lumen alter brain activity via receptor-mediated modulation of vagal afferent signaling.

**Figure 5:**
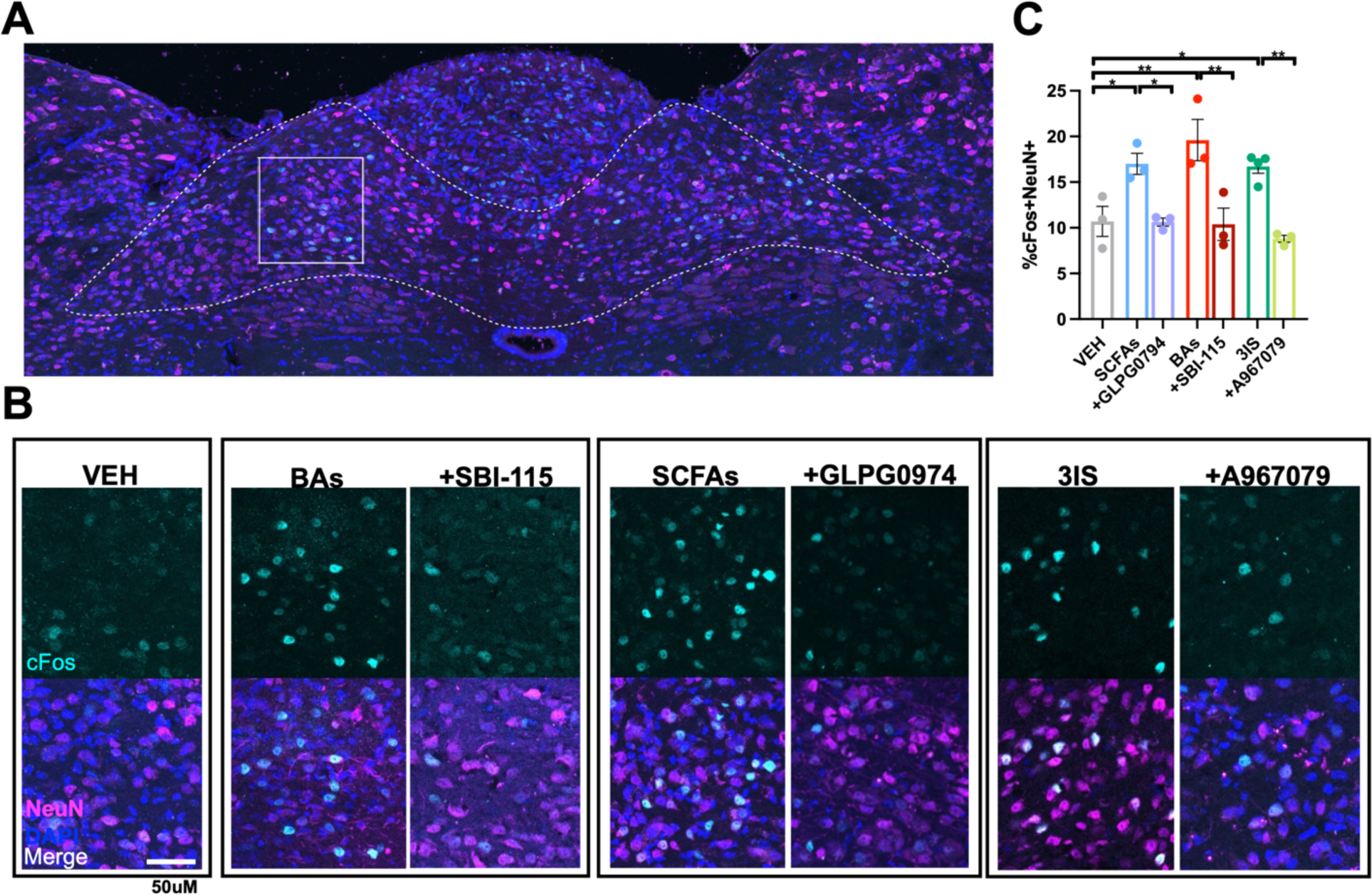
Microbial metabolites alter cFos levels in NTS neurons in a receptor-dependent manner. **A**) Representative image of full ROIs in mNTS for cFos quantification (dotted line) and inset images in B (solid line). **B-C**) Significant increase in the % of cFos+ NTS neurons following lumenal perfusion of BAs (cholate, 1240nM; glycocholate, 3.5nM; chenodeoxycholate, 42nM; alpha-muricholate, 142nM; beta-muricholate, 1080nM; deoxycholate, 390nM; taurodeoxycholate, 260nM; ursodeoxycholate, 74nM; taurohyodeoxycholate, 18.8nM; 7-ketodeoxycholate, 100nM; lithocholate, 390nM; taurolithocholate, 0.33nM, pooled, n = 3 mice), SCFAs (acetate 80uM, butyrate 22uM, propionate 10uM, pooled, n = 3 mice), and 3-IS (1uM, n = 4 mice) compared to vehicle (VEH, n = 3 mice) or corresponding receptor antagonists alongside each respective metabolite class (SBI-115, 200uM, n = 3 mice, GLPG0794, 10uM, n = 3 mice, A967079, 10uM, n = 3 mice). One-way ANOVA + Tukey. All data depicted as mean +/- SEM, *p < 0.05, **p < 0.01.

## DISCUSSION

Results from this study demonstrate that microbial colonization status, as well as acute manipulation of the gut microbiota and microbial metabolites, modulate vagal activity. In particular, we find that the gut microbiota regulates numerous small molecules in the small intestine and cecum. Moreover, administering select microbiome-dependent BAs, SCFAs, and 3IS at physiological concentrations and rates of peristalsis into the lumen of the small intestine stimulates vagal afferent neuronal activity. The vagal responses are elicited within relatively short timescales (<∼9 min) and are abrogated by pre- and co-administration of select receptor antagonists, suggesting active signaling between the gut microbiome and vagal afferents via excitatory metabolites.

The functional evidence provided in this study align with prior reports indicating that subdiaphragmatic vagotomy abrogates effects of microbial interventions on behaviors such as anxiety,^7^ depression,^57^ cognition,^60^ feeding,^46^ and social behaviors.^2,49^ Additionally, a few prior studies have reported that the gut microbiota and various microbial products regulate the transcriptome and excitability of vagal neurons.^58^ For instance, nodose neurons from mice reared GF exhibited altered gene expression profiles when compared to those from SPF mice,^61^ suggesting that the microbiome modulates the cellular state of vagal afferent neurons. In addition, bacterial supernatants from a cultured microbial community increased the intrinsic excitability of nodose neurons *in vitro*, through a mechanism that implicated a role for bacterial cysteine proteases.^62^ Another study reported that microbial single-stranded RNAs elevated vagal activity via Piezo1-mediated sensing by enterochromaffin cells.^23^ Moreover, the tryptophan metabolite indole has been reported to induce serotonin release onto colonic vagal afferents via TRPA1- mediated enteroendocrine cell activation.^43^ Together, these findings suggest that there exist multiple signaling factors and pathways by which the host-associated microbiota can impact vagal activity.

Aligning with the complexity of microbial influences on vagal activity, we observed that mice reared as GF exhibited significantly reduced afferent vagal nerve activity relative to mice reared with a conventional SPF microbiota, and while this effect was abrogated by colonizing GF mice with an SPF microbiota during adulthood, it was not fully recapitulated by depletion of the gut bacteria by oral ABX (**Figure 1C**). Many factors could contribute to this discrepancy. First, GF mice lack microbiota across all exposed body sites, whereas oral ABX treatment only partially ablates bacterial members of the oral and gastrointestinal microbiomes.^37^ As such, it is possible that the reductions in vagal tone seen in GF mice could be mediated by changes in both intestinal and non-intestinal vagal afferents and/or the presence of residual microbes or microbial products in the intestine of SPF mice treated twice daily with ABX by oral gavage. Moreover, the reported alterations in nodose gene expression in GF mice relative to SPF mice^61^ raise the question of whether there are early influences of microbiota status on vagal neuronal development, which are not captured by oral ABX treatment during adulthood. Further studies are needed to uncover the relative contributions of different vagal neuronal subtypes in mediating microbiome-induced alterations in vagal tone, as well as to what degree microbes endogenous to different parts of the gastrointestinal tract contribute to these alterations.

Despite the modest effects of oral ABX treatment in mice on reducing vagal tone (**Figure 1C**), we found that restricted perfusion of non-absorbable ABX through the lumen of the small intestine acutely reduced afferent vagal nerve activity. We additionally observed that afferent vagal nerve activity was restored by the re-introduction of SPF SI and cecal contents into the small intestine and that these increases in activity were driven, at least in part, by the small molecule fraction. This was not seen with lumenal perfusion of SI and cecal contents from GF mice, indicating a role for small molecules that are modulated by the gut microbiome. Notably, we did not observe an overt vagal nutrient response with perfusion of intestinal contents from GF mice, which could reflect the rapid digestion, absorption, and/or relatively low homeostatic concentration of microbiome-independent nutrients, such as glucose and sucrose, in the SI,^63^ as compared to the those exogenously delivered in other studies.^20-22^ Overall, these findings reveal that soluble microbiome-dependent metabolites from the lumen of the small intestine can acutely stimulate afferent vagal activity.

By untargeted metabolomic profiling of SI and cecal contents from conventionally colonized (SPF, CONV) and microbiota deficient (ABX, GF) mice, followed by *in vivo* screening of select microbial metabolites with or without pharmacological antagonists, we identified particular subclasses of microbiome-dependent molecules that activate vagal afferent neurons in a receptor-dependent manner upon administration to the small intestine. The metabolites—specific microbial BAs, SCFAs, and 3IS--promoted afferent vagal nerve activity with differing response kinetics, which could be due to differences in their physiological concentrations, rate of absorption, spatial localization of cognate receptors, and direct vs. indirect action, among other factors. Indeed, neuronal activation via GPCR signaling is dependent upon the concentration of the ligand, whereby differences can induce switching from G-protein coupled to G-protein independent signaling^64^ resulting in alterations in the downstream signal transduction pathways engaged during neuronal activation. In addition, microbiome-dependent BAs and SCFAs are reported to be absorbed by the intestinal epithelium, which offers the potential to activate gut-innervating vagal afferents through direct receptor binding,^15,16,18,19^ or through indirect interactions with diverse enteroendocrine cells, subsets of which can synapse directly onto vagal neurons.^65^ 3IS, however, has not been shown to be readily re-absorbed following secretion into the intestinal lumen, suggesting this metabolite likely acts through the indirect pathway in order to activate vagal afferents. It is also possible that select intestinal metabolites may access systemic circulation and act at extra-intestinal sites to modulate vagal activity, presumably with a time delay. In particular, lumenal microbiome-dependent BAs rapidly and transiently increased afferent vagal nerve activity, primarily via TGR5 signaling (**Figure 3A-C**). This may align with prior studies demonstrating that circulating BAs mediate release of cholecystokinin (CCK), and that CCK signaling is dynamic and rapidly desensitizes.^66^ In contrast, perfusion of SCFAs into the SI lumen increased afferent vagal nerve activity following a latency period in an FFAR2-dependent manner (**Figure 3D-F**), suggesting indirect activation of FFAR2-expressing epithelial cells and subsequent GLP-1 release.^67^ Finally, we found that microbiome-dependent 3IS in the small intestine elicited sustained afferent vagal nerve activity in a TRPA1-dependent manner (**Figure 3G-I**). This may align with a previous study wherein indole stimulated TRPA1+ colonic enteroendocrine cells to release serotonin and activate colon-innervating neurons.^43^ Our observations, considered together with existing vagal single cell transcriptomic data, raise the potential for both direct activation of afferent vagal neurons (via TGR5 and TRPA1) and indirect activation of epithelial cells or other cellular mediates (via FFAR2, TGR5, or TRPA1) by lumenal microbial metabolites. Future studies interrogating the cell-type specific role of metabolite receptors expressed on multiple cell types will aid in uncovering the precise differential effects of indirect versus direct signaling on vagal responses to microbial stimuli.

Recent studies demonstrate that lumenal nutrient cues, such as fats and carbohydrates, engage parallel vagal afferent neurons via labeled lines.^20-22^ However, further characterization of how different subclasses of diet- and microbiome-dependent small molecules engage gut-brain circuits involved in nutrient sensing is needed. We assessed the effects of acute lumenal perfusion of select microbial BAs, SCFAs, and 3IS (involved in dietary fat, carbohydrate, and protein metabolism, respectively) on afferent vagal neuronal calcium activity *in vivo*. We found that all three classes of microbial metabolites resulted in increased calcium activity in nodose neurons with varied kinetics (**Figure 4B**)-- BAs elicited a bimodal distribution of immediate vs. delayed responses, whereas SCFAs and 3IS mostly elicited delayed responses (**Figure 4D**), which aligns with the slow, gradual onset of afferent vagal nerve activity in response to SCFA and sustained onset of afferent nerve activity with 3IS perfusion. When assessing single-unit responses to sequential perfusion of two metabolite classes, microbiome-dependent BAs and SCFAs elicited calcium responses via largely non-overlapping subpopulations of afferent vagal neurons, whereas 3IS and BAs or SCFAs acted primarily via shared subpopulations (**Figure 4F**). These findings are supported by previous work demonstrating a shared role for both TGR5- and TRPA1-mediated alterations in digestion and satiety via epithelial CCK signaling,^31,68-71^ as well as TRPA1- and FFAR2-mediated alterations in host metabolism and feeding behaviors that have been reported to act via epithelial secretion of GLP-1.^67,72-74^ In contrast, previous work demonstrates that BA- and SCFA-mediated alterations in feeding behaviors act via distinct receptor-dependent signaling pathways.^49,75^ Future studies utilizing combinatorial functional strategies to elucidate the interplay between distinct metabolite effects on shared host behavioral and physiological processes will aid in uncovering the precise cellular crosstalk that may mediate our observed results.

Despite evidence for vagal chemosensory pathways mediating communication from the intestinal lumen to the brain,^22,58,76^ effects of specific microbial metabolites and their associated receptors on CNS neuronal activity remain unclear. We therefore addressed the effects of lumenal perfusion of BAs, SCFAs, and 3IS alongside their respective receptor antagonists for TGR5, FFAR2, and TRPA1 on medial NTS neuronal activation by immunofluorescence detection of the immediate early gene cFos. We found that all metabolite classes significantly increased NTS neuronal activation, which could be prevented by pre-and co-perfusion of antagonists (**Figure 5**). Together, these data suggest that lumenal metabolites activate CNS neurons via receptor-dependent vagal signaling and may have downstream effects on CNS responses that contribute to behavior. However, more work is needed to determine the downstream targets and differential effects of these gut-to brain signaling pathways on CNS physiology.

Findings from this study highlight lumenal microbial metabolites derived from various sources of dietary macronutrients--fats (BAs), complex carbohydrates (SCFAs), and proteins (3IS)—that differentially activate vagal afferent neurons via receptor-mediated signaling in order to convey information to the brain. Following the ingestion of dietary fats, BAs are released from the liver into the SI lumen where they undergo chemical transformations, such as deconjugation and dehydroxylation, which are carried out by gut microbes.^77^ Enzymes capable of catalyzing such reactions are found across all major bacterial phyla,^78^ suggesting a broad role for the microbiota in regulating the lumenal BA pool. Similarly, SCFAs are derived from microbial metabolism of dietary fibers that are otherwise non-digestible by the host, with differential production by bacterial members of the phyla *Bacteroidetes* (*Bacteriodata;* acetate, propionate) and *Firmicutes* (*Bacillota*; butyrate).^79,80^ Tryptophan derivatives, such as indole, are produced by pathobionts, such as *Escherichia coli*, *Enterococcus faecalis,* and *Edwardsiella tarda,^81^* before being hydroxylated and sulfated in the liver and secreted as 3IS into the small intestine. As levels of dietary metabolites and microbiome composition can fluctuate depending on meal time^82,83^, further examination into circadian effects of diet- and microbiome-dependent metabolites on vagal afferent neuronal activity is warranted.

Understanding the temporal variation in the bioavailability of neuromodulatory microbial metabolites and in vagal activity could reveal important insights into the functional role of vagal interoception of different types of microbial metabolites. As proof of principle, vagal TGR5 signaling mediated the anorexigenic effects of circulating BAs,^31^ whereas vagal FFAR3 signaling mediated the effects of circulating SCFAs on satiety.^49^ However, further studies on whether the vagal circuits engaged by lumenal microbial metabolites modulate analogous or additional CNS-associated behaviors remains to be determined. Circuit tracing studies have uncovered polysynaptic connections from gut-innervating vagal afferent neurons to higher order brain regions such as substantia nigra and hippocampus,^84^ suggesting the potential for microbial metabolites to impact complex behavioral responses beyond those involved in feeding. BAs, SCFAs, and 3IS have been associated with alterations in behavioral endophenotypes of anxiety and depression,^52,85,86^ cognitive impairment,^87^ and motor deficits,^3,86^ each which has been linked to vagal signaling and alterations in the gut microbiota.^3,9,10^ Overall, more research is needed to determine brain and behavioral responses to vagal interoception of lumenal microbial metabolites, and to further evaluate the potential to leverage the microbiome to modify neuronal signaling across the gut-brain axis.

## ACKNOWLEDGEMENTS

We thank members of the Hsiao laboratory for their guidance and review of the manuscript; members of the UCLA Goodman Luskin Microbiome Center Gnotobiotics Core Facility for technical support; Dr. Stephen Liberles for critical training on intestinal perfusion and vagal nerve electrophysiology, Dr. Daniel Aharoni for helpful advice regarding calcium imaging data analysis; Dr. Scott Kanoski and Dr. Diego Bohorquez for helpful feedback on the project; and Dr. Baljit Khakh for allowing usage of his osmometer. This work was supported by funds from an NIH Ruth L. Kirschstein National Research Service Awards (F31 NS118966 and T32 GM007185), UCLA Hyde Pre-doctoral Fellowship, and UCLA Dissertation Year Fellowship to K.G.J., UCLA Hyde Pre-doctoral Fellowship to S.A.K., American Heart Association Fellowship to C.S., and NINDS grant (R01 NS115537) to E.Y.H. E.Y.H. is a New York Stem Cell Foundation – Robertson Investigator. This research was supported in part by the New York Stem Cell Foundation. This project has been made possible in part by grant number 2018-191860 from the Chan Zuckerberg Initiative DAF, an advised fund of Silicon Valley Community Foundation.

## AUTHOR CONTRIBUTIONS

K.G.J., S.A.K., C.S., D.M., and E.L. performed the experiments and analyzed the data, H.E.V., J.P., L.Y., S.C.M., F.E.S. provided key technical guidance and resources. K.G.J. and E.Y.H. designed the study, K.G.J. and E.Y.H. wrote the manuscript. All authors discussed the results and commented on the manuscript.

## DECLARATION OF INTERESTS

The authors declare no competing interests.

## DIVERSITY AND INCLUSION

One or more of the authors of this paper self-identifies as an underrepresented ethnic minority in science. While citing references scientifically relevant for this work, we also actively worked to promote gender balance in our reference list.

## STAR★METHODS

Detailed methods are provided and include the following:

- KEY RESOURCES TABLE
- CONTACT FOR REAGENT AND RESOURCE SHARING
- EXPERIMENTAL MODELS AND SUBJECT DETAILS

- Mice
- METHOD DETAILS

- Antibiotic Treatment and Conventionalization
- Preparation of Antibiotics, Small-Intestinal, and Cecal Contents
- *In Vivo* Vagal Electrophysiology
- Metabolomics
- Preparation of Metabolite Pools and Single Metabolites
- *In Vivo* Calcium Imaging
- Immunohistochemistry for cFos Detection
- QUANTIFICATION AND STATISTICAL ANALYSIS
- DATA AND SOFTWARE AVAILABILITY

## KEY RESOURCES TABLE

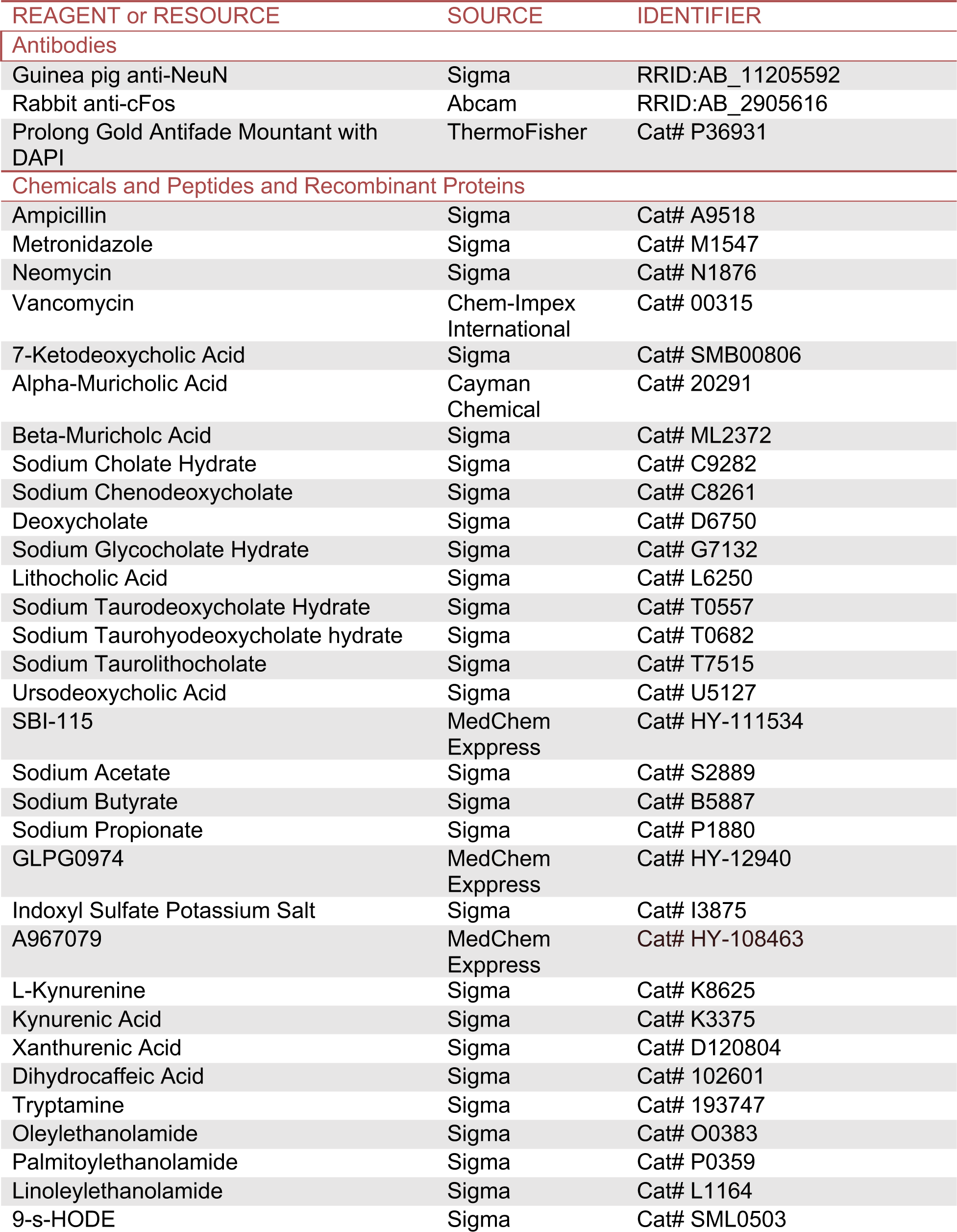
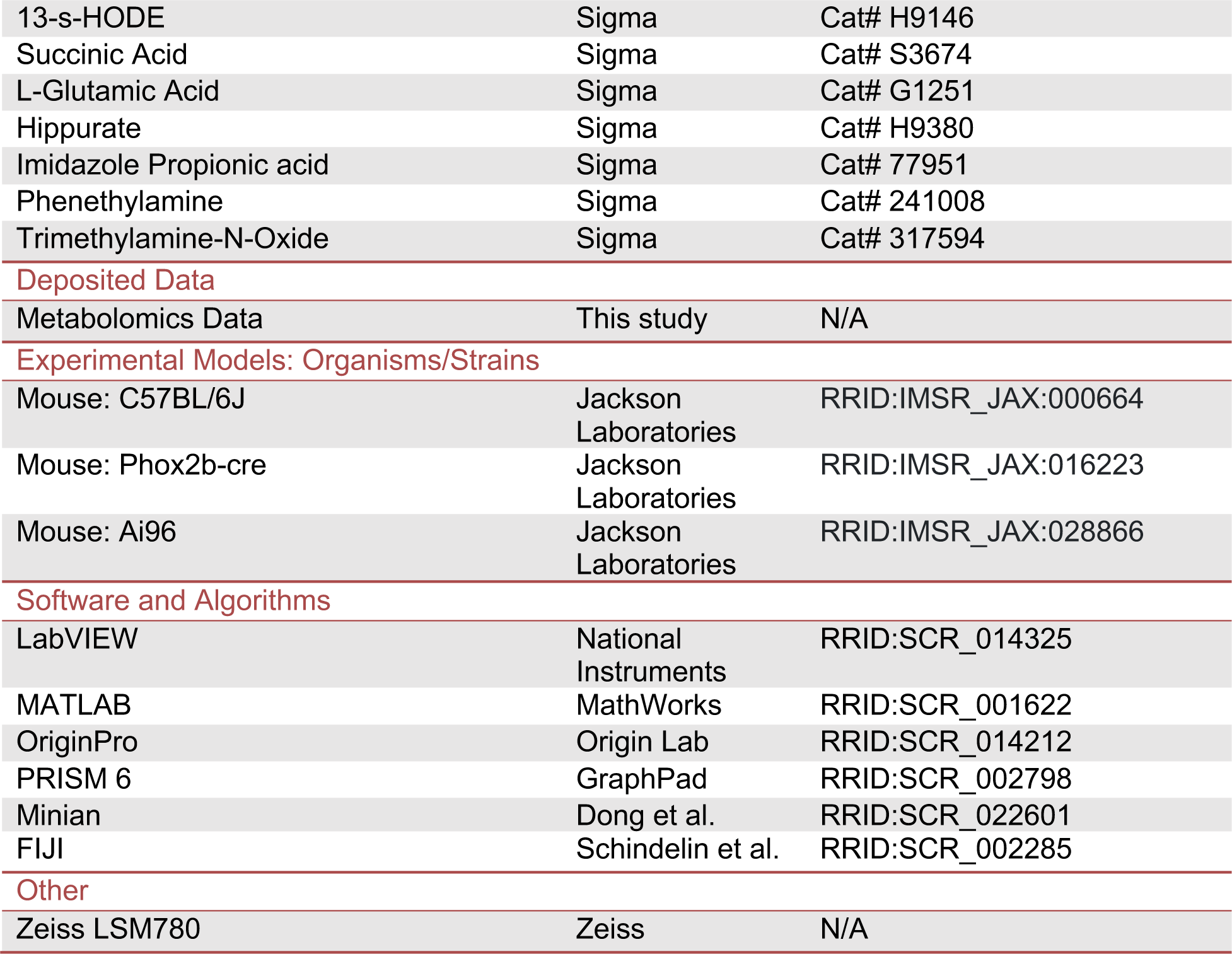

## RESOURCE AVAILABILITY

### Lead Contact

Further information and requests for resources and reagents should be directed to and will be fulfilled by the Lead Contact, Elaine Hsiao

### Materials Availability

This study did not generate new unique reagents.

### Data and Code Availability

All source data are included as supplementary tables in this manuscript.

## EXPERIMENTAL MODELS AND SUBJECT DETAILS

### Mice

All experimental procedures were carried out in accordance with US NIH guidelines for the care and use of laboratory animals and approved by the UCLA Institutional Animal Care and Use Committees. Mice used for data collection were males, at least 6 weeks of age. C57BL/6J mice were purchased from Jackson laboratories (stock no. 000664), reared as SPF or rederived as GF and bred in flexible film isolators at the UCLA Center for Health Sciences Barrier Facility. Ai96 (JAX stock no. 028866), *Phox2b-cre* (JAX stock no.016223), were obtained from Jackson laboratories and bred at the UCLA Biomedical Sciences Research Building barrier facility. Mice were housed on a 12-h light-dark schedule in a temperature-controlled (22-25°C) and humidity-controlled environment with *ad libitum* access to water and standard chow (Lab Diet 5010).

## METHOD DETAILS

### Antibiotic treatment and conventionalization

Adult SPF mice were gavaged twice daily for 1 week with a cocktail of vancomycin (50mg/kg), neomycin (100mg/kg) and metronidazole (100mg/kg) every 12 hours daily for 7 days. Ampicillin (1mg/mL) was provided *ad libitum* in drinking water. For conventionalization, fecal samples were collected from adult SPF C57BL/6J mice and homogenized in 1mL pre-reduced PBS per pellet. 100uL of the homogenate was administered by oral gavage to recipient GF mice.

### Preparation of antibiotics, small-intestinal, and cecal contents

Vancomycin and neomycin were diluted in water to a final concentration of 1mg/mL for all antibiotic perfusion experiments, then sterile filtered. Vehicle for all antibiotic perfusion experiments was water. For SPF SI and cecal content preparations adult SPF male mice were euthanized, and SI and cecal lumenal contents were snap-frozen in liquid nitrogen. Equal weights of frozen SI and cecal content were then combined and diluted to a concentration of 0.1g/mL wet weight in sterile-filtered PBS. Samples were then centrifuged at 500g for 5 minutes to pellet out any large dietary components, and supernatants were used for lumenal perfusion. GF SI/cecal contents were collected in the same way from donor GF adult male mice. Sterile-filtered SI/cecal contents were prepared by vacuum-filtering SPF SI/cecal content supernatants through a 0.2μm filter. Sterile-filtered PBS was used as vehicle in all SI/cecal perfusion experiments.

### *In vivo* vagal electrophysiology

Baseline vagal tone recordings: Adult SPF, GF, ABX, or CONV male mice were anesthetized with isoflurane (5%) and maintained at 1.8% throughout the experiment. The cervical vagus nerve was exposed, transected inferior to the nodose ganglion and placed across two platinum iridium wires (insulated except for a short segment in contact with the nerve) for recording of baseline vagal tone. Recordings were conducted for 10 minutes. Vagal recordings with lumenal perfusion: the cervical vagus nerve was prepared as above in adult male SPF mice. Additionally, a 20-gauge gavage needle attached to a peristaltic pump (Cole Parmer) with separate tubing for each infusion solution was inserted into the duodenal lumen and secured with sutures. An outflow port was generated by transecting the small intestine ∼10cm distal to the inflow site. During recording, vehicle was first perfused through the lumen at a constant rate for 10 minutes to establish a baseline following surgery at a flow rate of 250 ul/minute.^20^ Following the baseline period, stimuli were introduced into the small-intestinal lumen and perfused at the same rate for the remainder of the experiment. Data Acquisition: a differential amplifier was used (A-M Systems LLC). The gain was set to 1000x and a bandpass filter was applied (300Hz-5kHz). The signal was digitized at 20kHz using a data acquisition board (National Instruments) under the control of LabView software Data Analysis: Spikes were detected using an SO-CFAR threshold (window duration of 1501/8000, guard duration 10/8000) to generate an adaptive threshold 4SD above RMS noise^88^. Firing rates were calculated by generating 10s (baseline vagal tone) or 30s (perfusion experiments) bins and then applying a Savitszky-Golay filter (OriginPro) of 10 points. Baseline vagal tone was defined as the average of the final 300s of recording. All raw values were normalized to the SPF cohort average for the rig on which the recordings took place. Perfusion experiments: baseline values were defined as the average frequency of the final 60s of recording in the initial baseline period (F_0_). Frequency of the recording was then normalized to the baseline value within-subject (F/F_o_). Area under the curve was calculated for each stimulus window and defined as the integral of frequencies over the stimulus window.

### Metabolomics

Samples were collected from adult SPF, GF, ABX, or CONV mice. Lumenal contents were collected from the first 3cm of small intestine and the entirety of the cecum, then snap frozen in liquid nitrogen and stored at -80°C. Samples were prepared using the automated MicroLab STAR system (Hamilton Company) and analyzed on gas chromatography (GC)-mass spectrometry (MS), liquid chromatography (LC)-MS and LC-MS/MS platforms by Metabolon, inc. Protein fractions were removed by serial extractions with organic aqueous solvents, concentrated using a TurboVap system (Zymark) and vacuum dried. For LC/MS and LC-MS/MS, samples were reconstituted in acidic or basic LC-compatible solvents containing > 11 injection standards and run on a Waters ACQUITY UPLC and Thermo-Finnigan LTQ mass spectrometer, with a linear ion-trap front-end and a Fourier transform ion cyclotron resonance mass spectrometer back-end. For GC/MS, samples were derivatized under dried nitrogen using bistrimethyl-silyl-trifluoroacetamide and analyzed on a Thermo-Finnigan Trace DSQ fast-scanning single-quadrupole mass spectrometer using electron impact ionization. Chemical entities were identified by comparison to metabolomic library entries of purified standards. Following log transformation and imputation with minimum observed values for each compound, data were analyzed using one-way ANOVA to test for group effects. P and q-values were calculated based on two-way ANOVA contrasts. Principal components analysis was used to visualize variance distributions. Supervised Random Forest analysis was conducted to identify metabolomics prediction accuracies.

### Preparation of metabolite pools, single metabolites, and receptor antagonists

All working metabolite solutions were made up in PBS and up to 0.05% DMSO, brought to a pH of 7.3, and sterile filtered. The concentrations for each metabolite pool, individual metabolites, and receptor antagonists were determined from serum metabolomics data in the lab (data not shown) and existing literature and are as follows: Tryptophan metabolites (kynurenine 100uM,^89^ kynurenic acid 16.1uM,^90^ xanthurenate 1uM,^91^ dihydrocaffeate 30nm, tryptamine 0.03uM, pooled), FAEs (oleylethanolamide (OEA), palmitoylethanolamide (PEA), linoleylethanolamide (LEA) 10uM, pooled)^92^, HODEs (9-s- and 13-s-HODE,1uM, pooled), succinate (2mM), glutamate (50mM), bile acids (BAs, cholate 1240nM, glycocholate 3.5nM, chenodeoxycholate 42nM, alpha-muricholate 142nM, beta-muricholate 1080nM, deoxycholate 390nM, taurodeoxycholate 260nM, ursodeoxycholate 74nM, taurohyodeoxycholate 18.8nM, 7-ketodeoxycholate 100nM, lithocholate 390nM, taurolithocholate .33nM, pooled), m-tolyl 5-chloro-2-[ethylsulonyl] pyrimidine-4-carboxylate (SBI-115, 200uM)^48^, short-chain fatty acids (SCFAs, acetate 80uM, butyrate 22uM, propionate 10uM, pooled), 4-[[(R)-1-(benzo[b]thiophene-3-carbonyl)-2-methyl-azetidine-2-carbonyl]-(3-chloro-benzyl)-amino]-butyric acid 99 (GLPG0974, 10uM)^93^, hippurate (2uM)^94^, Trimethylamine-N-oxide (TMAO, 3uM)^95^, imidazole propionate (200nM),^96^ phenethylamine (PEA, 100uM)^13^, 3-indoxyl sulfate (3IS, 1uM)^97^, or (1E,3E)-1-(4-Fluorophenyl)-2-methyl-1-penten-3-one oxime (A967079, 10uM).^44^ Vehicle for tryptophan metabolites, FAEs, MFAs, succinate, glutamate, SCFAs, TMAO, hippurate, imidazole propionate, and phenethylamine was sterile-filtered PBS. Vehicle for BAs was 0.05% DMSO in PBS, Vehicle for 3-IS was 1uM KCl in PBS.

### *In vivo* calcium imaging

Vagal afferent neuron imaging was performed as previously described.^20^ In brief, SPF mice were anesthetized with a cocktail of ketamine and xylazine (100 mg/kg and 10 mg/kg, intraperitoneal), tracheotomized, and maintained on 1.5% isoflurane for the remainder of the surgery and imaging session via ventilator (Kent Scientific). Body temperature was maintained at 37°C using an electrical heating pad and rectal temperature probe (Kent Scientific). The vagus nerve was transected superior to the jugular ganglion and vagal ganglia were then embedded between two 5mm-diameter glass coverslips (neuVitro) with silicone adhesive (KWIK-SIL, World Precision Instruments). Imaging was conducted on a Zeiss LSM 780 confocal microscope at a frame rate of 1Hz. Analysis of imaging data: Imaging data were analyzed using custom Python scripts based on the Minian pipeline.^98^ Imaging data was registered to collect for motion artifacts and denoised using a median filter. A constrained non-negative matrix factorization (CNMF) algorithm was used to identify single cells and extract calcium activity. CNMF output regions of interest were then manually inspected to remove non-neuronal signals. Metabolite stimuli order was randomized to account for order-specific effects on neuronal activity. For single-unit analysis in response to sequential metabolite application, neuronal regions of interest were cross-registered across stimulus runs and compared to determine single- or dual-responsivity to metabolite classes. The baseline period for the calcium signal was calculated as the average of the last 120 seconds before stimulus onset (F_o_). Responses were reported in units of baseline fluorescence. Cells were considered responsive to stimuli if the maximal ΔF/F signal following stimulus onset was i) greater than 2 standard deviations above the baseline fluorescence, and ii) the mean ΔF/F signal over a 20s window around peak response was >50% of the baseline value. Only units that displayed a ΔF/F signal >4 standard deviations over the baseline fluorescence in response to an electrical stimulus were included in analysis. For visualization, metabolite-responding units were hierarchically clustered over the entire time-course of each experiment.

### Immunohistochemistry for cFos

Lumenal perfusion of stimuli: Mice were anesthetized with 5% isoflurane and maintained at 2% for the entirety of the surgery. To begin intralumenal perfusion, the small intestine was transected at the pyloric sphincter, the junction between the stomach and duodenum, to create an inflow port. Subsequently, a gavage needle attached to a peristaltic pump was inserted into the duodenal lumen. An outflow port was made by transecting the small intestine 3 centimeters distal to the inflow port. All lumenal contents were flushed from the small intestine with PBS. Animals were then continually perfused with PBS at 250uL/min flow rate for a 10-minute baseline period, followed by perfusion of stimuli for 30 minutes at a constant flow rate to account for any mechanical distension. Following stimulus perfusion, mice remained under anesthesia for one hour to allow for cFos induction before tissue harvesting. Following the one-hour rest period, animals were sacrificed, and tissues were harvested and fixed via intracardial perfusion of ice-cold PBS followed by 4% paraformaldehyde (PFA). Brains were then post-fixed in 4% PFA at 4°C for 3 hours followed by an overnight incubation in 30% sucrose at 4°C for cryoprotection. Ganglia were positioned bulb-side down in optimal cutting temperature compound (OCT compound) and frozen for cryostat sectioning. Tissues were then sectioned at 30µm and mounted on a microscope slide for immunohistochemical processing (IHC). Slides were thawed for 10 minutes at room temperature in a humidified chamber to prevent the tissue from drying and then permeabilized in 0.5% Triton/0.05% tween-20 in PBS (PBS-TT). Blocking solution consisting of 5% normal goat serum (NGS) in PBS-TT was applied to the tissue and allowed to incubate at room temperature for two hours to prevent non-specific antibody binding and reduce background staining. The tissue sections were then incubated overnight at 4°C with primary antibodies (rabbit anti-cFos 1:500, and guinea pig anti-NeuN, 1:500) in blocking solution (5% NGS + PBS-TT). The following day, slides were washed three times, five minutes per wash, with PBS-TT before incubating at room temperature for two hours with secondary antibodies (goat anti-rabbit 488 1:1000, goat anti-guinea pig 568 1:1000, and DAPI 1:1000) in blocking solution (5% NGS + PBS-TT). Confocal Imaging and Quantification Analysis: Images were obtained using a 20x air objective (NA 0.8) on an upright Zeiss LSM 780 confocal microscope. Z-stacks were acquired for three technical replicates of NTS brain tissue and maximum-intensity projections were generated for subsequent analysis in ImageJ. NTS sections were selected and ROI’s were drawn based off of the Allen Mouse Brain Atlas. First, NeuN positive (NeuN+) cells that were each confirmed to colocalize with a DAPI nucleus were counted using the multi-point tool to obtain the total number of neurons. Subsequently, cFos+ cells were counted using the multi-point tool by confirming colocalization of a cFos immunofluorescence signal with NeuN and DAPI. For each image, the total number of cFos+ neurons were divided by the total number of NeuN+ cells to obtain the percentage of cFos+ neurons. Finally, the percentage of cFos+ neurons for all technical replicates of NTS slices per animal were averaged to obtain a biological n=1 and find the overall percentage of cFos+ neurons.

## QUANTIFICATION AND STATISTICAL ANALYSIS

Statistical analysis was performed using Prism software version 8.2.1 (GraphPad). Data were assessed for normal distribution and plotted in the figures as mean ± SEM. For each figure, *n* = the number of independent biological replicates. Outliers were identified with ROUT using a threshold of q = 2% for all electrophysiological recordings. Differences between two treatment groups were assessed using two-tailed, unpaired Student *t* test with Welch’s correction. Differences among >2 groups with only one variable were assessed using one-way ANOVA. If groups were determined to have significantly different variances, groups were assessed with Brown-Forsythe and Welch ANOVA + Games-Howell testing or Wilcoxon matched-pairs signed rank test. Significant differences emerging from the above tests are indicated in the figures by *p < 0.05, **p < 0.01, ***p < 0.001, ****p < 0.0001. Notable non-significant (and non-near significant) differences are indicated in the figures by “n.s.”.

## SUPPLEMENTAL INFORMATION

Supplemental Information includes 2 figures, 2 tables and all source data, and can be found with this article.

**Supplemental Figure 1:**
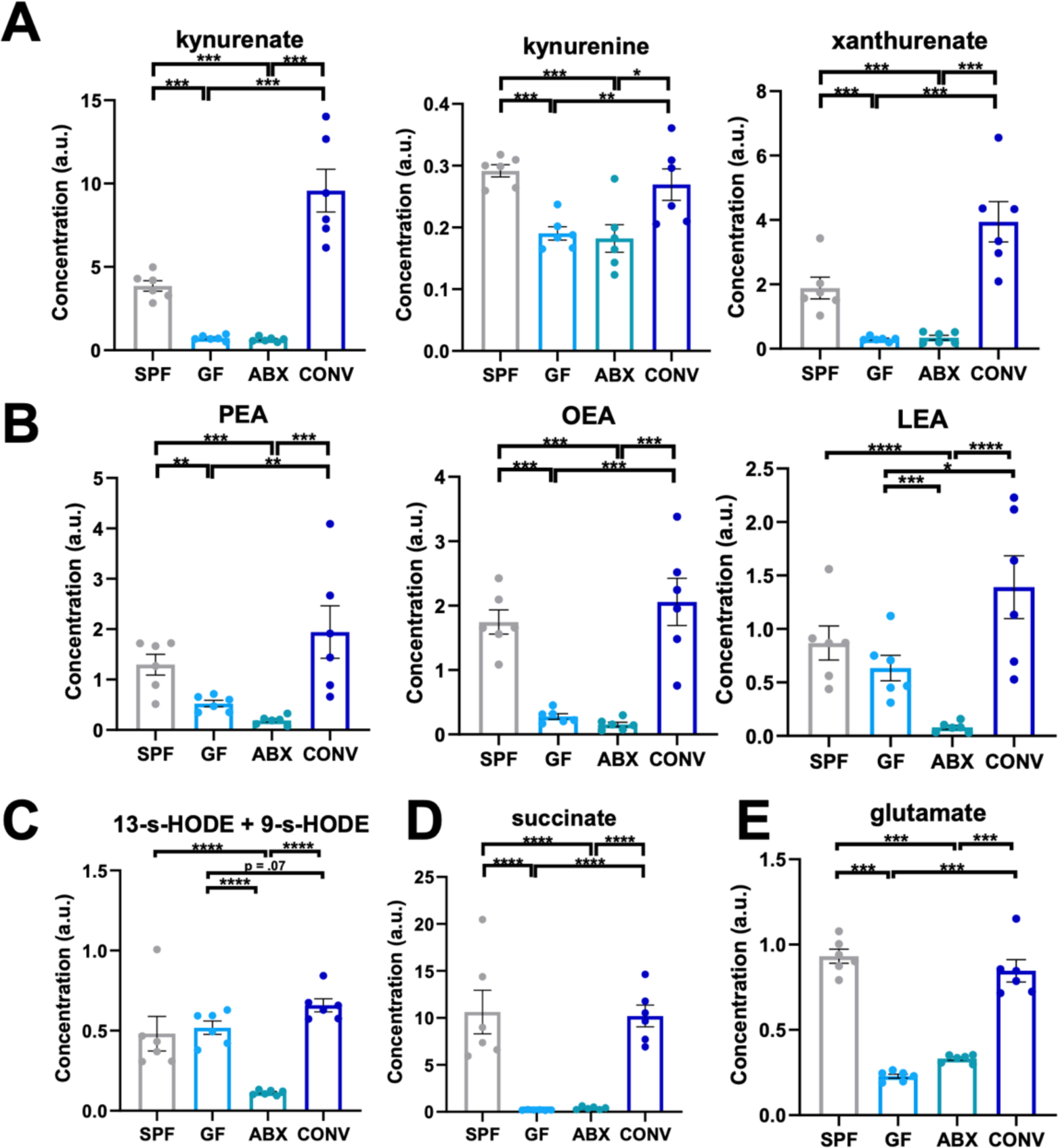
Microbially-modulated cecal metabolites that were screened for effects on afferent vagal nerve activity. Lumenal levels of microbially modulated **A**) tryptophan metabolites, **B**) Fatty-acid ethanolamides (FAEs) PEA (palmitoyl ethanolmaide), OEA (oleoyl ethanolamide), LEA (linoleoyl ethanolamide), **C**) monohydroxy fatty acids (MFAs) 9-s-HODE and 13-s-HODE, **D**) succinate, **E**) glutamate. Welch’s t-test. All data displayed as mean +/- SEM, *p < 0.05, **p < 0.01, ***p < 0.001, ****p<.0001.

**Supplemental Figure 2:**
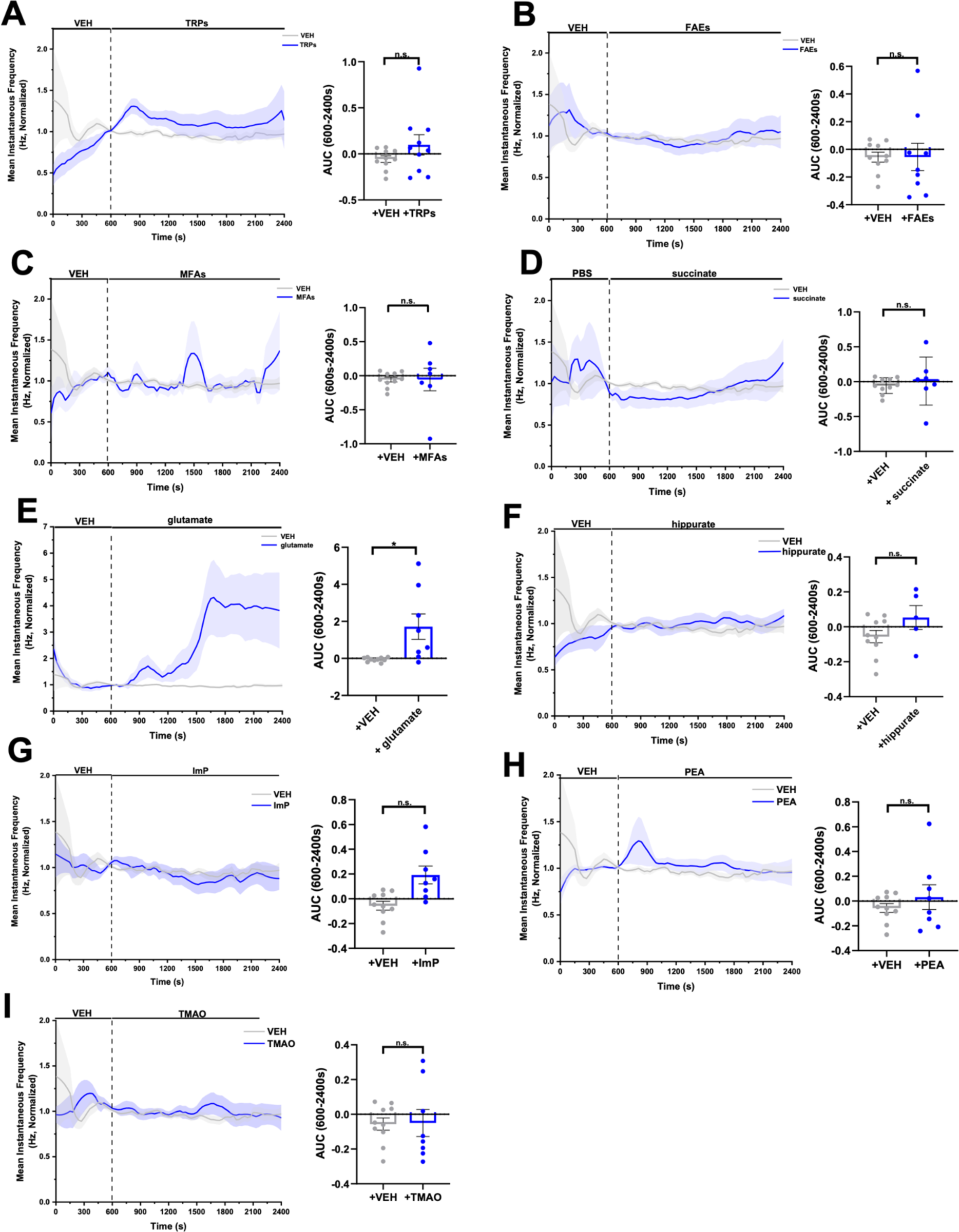
Lumenal perfusion of select microbially-modulated metabolites does not alter vagal afferent nerve activity *in vivo*. Afferent vagal nerve firing rate in response to lumenal perfusion of VEH (PBS, n = 10 mice) or **A**) tryptophan metabolites (TRPs) (kynurenine 100uM, kynurenic acid 16.1uM, xanthurenate 1uM, dihydrocaffeate 30nm, tryptamine 0.03uM, pooled, n = 10 mice), **B**) fatty-acid ethanolamides (FAEs, oleylethanolamide (OEA), palmitoylethanolamide (PEA), linoleylethanolamide (LEA), 10uM, pooled, n = 9 mice), **C**) monohydroxy fatty acids (MFAs) (9-s- and 13-s-HODE,1uM, pooled, n = 7 mice), **D**) succinate (2mM, n = 7 mice), **E**) glutamate (50mM, n = 8 mice), **F**) hippurate (2uM, n = 5 mice), **G**) imidazole propionate (ImP, 200nM, n = 8 mice),**H**) phenethylamine (PEA, 100uM, n = 8 mice), or **I**) trimethylamine-N-oxide (TMAO, 3uM, n= 8 mice). Welch’s t-test for each comparison. All data displayed as mean +/- SEM, *p < 0.05, **p < 0.01, ***p < 0.001

## Notes

### Competing Interest Statement

The authors have declared no competing interest.

